# Effects of HA1 (a Probucol Analogue) and ApoC3 siRNA on Lipoprotein-Amyloid Metabolism, Neurovascular Integrity and Cognitive Function in db/db Mice

**DOI:** 10.1101/2024.11.05.622070

**Authors:** Arazu Sharif, John Mamo, Virginie Lam, Gerald F Watts, Giuseppe Luna, Michael Nesbit, Hani Al-Salami, Armin Mooranian, Ryu Takechi

**Affiliations:** Curtin Health Innovation Research Institute, Faculty of Health Sciences, Curtin University, Perth, WA, Australia; Perron Institute for Neurological and Translational Research, Perth, WA, Australia; Curtin Medical School, Faculty of Health Sciences, Curtin University, Perth, WA, Australia; School of Public Health, Faculty of Health Sciences, Curtin University, Perth, WA, Australia; School of Medicine,, University of Western Australia, Perth, WA, Australia; Department of Cardiology, Royal Perth Hospital, Perth, WA, Australia; School of Pharmacy, University of Otago, Dunedin, New Zealand

**Keywords:** Type 2 Diabetes, Alzheimer’s Disease, Lipoprotein-Aβ, Blood-Brain Barrier, Probucol, ApoC3 siRNA, Anxiety-Like Behavior

## Abstract

**Background and aims:** Chronically elevated levels of circulating lipoprotein-amyloid-β (Aβ) are implicated in the disruption of the blood-brain barrier and the initiation of a neurodegenerative cascade leading to Alzheimer’s disease (AD). Type 2 diabetes is associated with BBB dysfunction, dyslipidaemia, and an increased risk of AD. However, alterations in triglyceride-rich lipoproteins (TRL)-Aβ homeostasis and its downstream effects on the BBB in a diabetes context remain explored. This study aimed to 1) investigate, in a preclinical model of diabetes, the hypothesis that diabetes-induced impairments in TRL-Aβ metabolism might compromise BBB integrity and exacerbate cognitive function and behavioural changes, and 2) assess the efficacy of interventions that improve TRL catabolism, including probucol, HA1, and ApoC3 siRNA, to prevent disease progression by lowering circulating TRL-Aβ levels.

**Methods:** Five-week-old db/db mice underwent a 9- or 23-weeks dietary interventions with probucol, a probucol prodrug HA1, or a standard diet with four-weekly injections of ApoC3 siRNA. Db/+ mice served as negative controls for each treatment duration. Blood levels of Aβ and ApoB were measured using ELISA. Immunofluorescence imaging was used to quantify enterocytic levels of Aβ and ApoB, and assess changes in neurovascular integrity (IgG, PDGFRβ, ZO1, occludin), neuroinflammation (GFAP, Iba1), and cerebral oxidative stress (8OHdG).

**Results:** Our results indicate that diabetes increased the abundance of plasma amyloid, specifically Aβ42, which correlated with enterocytic abundance, suggesting exaggerated postprandial excretion. Disrupted plasma amyloid homeostasis was associated with BBB breakdown, including diminished barrier function, and the loss of pericytes and astrocytes. Provision of the probucol analogue, HA1, normalised plasma and enterocytic amyloidemia concomitant with the preservation of the neurovascular junction. Treatment with ApoC3 siRNA attenuated plasma Aβ42 and modestly reduced neurovascular inflammation.

**Conclusion:** The findings further support the hypothesis that aberrant peripheral metabolism of lipoprotein-Aβ is associated with microvascular corruption and the development of anxiety-like behaviour. HA1 is more effective than probucol or ApoC3 siRNA in positively modulating lipoprotein-amyloid homeostasis in db/db mice and maintaining central capillary integrity.

**Highlights:**

- Enterocytic and plasma lipoprotein-Aβ levels are increased in diabetic db/db mice.
- Elevated plasma lipoprotein-Aβ levels correlate with BBB breakdown and anxiety-like behavior.
- HA1 lowers enterocytic and plasma Aβ levels, protects BBB, alleviates oxidative stress and anxiety.
- ApoC3 siRNA lowers plasma lipoprotein-Aβ, mitigates neuroinflammation and anxiety.

## 1. Introduction

Alzheimer’s disease (AD) is the most common form of dementia, marked by progressive cognitive decline, neuronal loss and neuroinflammation. Increasing evidence suggests that diabetes mellitus (DM), particularly type 2 diabetes (T2D), is a major risk factor for the development of AD and other neurodegenerative conditions [1]. This link is supported by the observation that diabetes is associated with brain atrophy, blood-brain barrier (BBB) dysfunction and the deposition of amyloid-beta (Aβ), a hallmark of AD pathology [2]. However, the mechanisms by which diabetes contributes to AD pathogenesis remain incompletely understood.

Recent studies highlight the peripheral metabolism of lipoprotein-bound Aβ, particularly within triglyceride-rich lipoproteins (TRLs), as a key contributor to the neurovascular pathology seen in AD [3]. Preclinical models have shown that diets high in fat can exacerbate the secretion of TRL-bound Aβ, leading to BBB disruption and the extravasation of this lipoprotein-Aβ complex into the brain parenchyma [4–7]. Once in brain, these complex trigger neurovascular inflammation, neuronal loss and cognitive deficits [3,5]. Particularly compelling evidence has come from genetically engineered mice that synthesize human Aβ exclusively in the liver, showing a clear peripheral contribution of Aβ to neurodegeneration [8].

Previous mouse models investigating high-fat diet-induced diabetes are consistent with a lipoprotein-amyloid and capillary axis for disease, but clearly confounded by the independent effects of dietary fat on the metabolic and neurovascular systems. These models do not fully capture the impact of insulin resistance—an underlying feature of type 2 diabetes that plays a critical role in both systemic and central metabolic dysfunction. To model the diabetes-AD connection more accurately, it is essential to consider mouse strains that better replicate insulin resistance-induced diabetes, independent of high-fat diet interventions.

In the context of diabetes, insulin resistance may promote excessive secretion of TRL-Aβ, exacerbated by metabolic disruptions such as hyperphagia, hypertrophy, obesity and dysregulated hepatic lipid metabolism. This “endocrine axis” of TRL-Aβ secretion in insulin resistance and diabetes may contribute to the development of AD by increasing peripheral Aβ burden. Studies using rodent models of insulin resistance-induced diabetes (without high-fat diet interventions) would provide a clearer picture of how this lipoprotein-Aβ/capillary axis drives BBB breakdown, neuroinflammation, and cognitive decline. Impaired hepatic clearance of TRL-Aβ and prolonged vascular exposure we postulate would allow these complexes to accumulate at the BBB, where they compromise barrier function. Considering these putative physiological aberrations in peripheral metabolism of lipoprotein-Aβ may offer a novel therapeutic approach to mitigating AD progression in diabetic patients.

Probucol, a lipid-lowering agent, has shown promise in reducing circulating TRL-Aβ levels by suppressing secretion of amyloid with nascent TRL, promoting clearance of TRL-Aβ and protecting BBB integrity in models of metabolic dysfunction, including high-fat diet-induced diabetes [9–11]. However, its clinical utility is limited by its poor bioavailability[12]. HA1, a novel probucol analogue conjugated with lithocholic acid, has demonstrated improved pharmacokinetics and may have greater therapeutic potential. ApoC3, a key regulator of TRL catabolism, is elevated in diabetes [13] and impairs TRL clearance by inhibiting lipoprotein lipase (LPL)-dependent triglyceride hydrolysis and receptor mediated hepatic clearance of the remnant lipoprotein thereafter [14], pontentially increasing vascular exposure to TRL-Aβ.

In this study, we aim to explore the role of TRL-Aβ in insulin resistance-induced BBB dysfunction and neurovascular inflammation utilizing leptin-receptor deficient db/db mice. We will also investigate whether interventions with native probucol, HA1 or ApoC3 siRNA can normalise TRL-Aβ metabolism, protect BBB integrity and prevent neuroinflammation and cognitive decline.

## 2. Materials and methods

### 2.1. Animals and interventional design

All procedures were approved by the Curtin Animal Ethics Committee (ARE 2020-12) and complied with the Australian Code for the Care and Use of Animals, 8th Edition. Five-week-old male db/db and db/+ C57BLK/6J mice were obtained from Jackson Laboratory and bred at the Animal Resource Centre (WA). Mice were housed with controlled air pressure, 12:12 light/dark cycles, and temperature at Curtin University’s Life Science Research Facility. After one week of acclimation, mice were divided into two age groups: 14 and 28 weeks. Within each group, 12–14 db/db mice were randomly assigned to standard chow (AIN93M, Speciality Feeds, WA) or one of three treatments: (1) chow supplemented with 1% probucol (Beijing Natural-Med Biotechnology), (2) chow with 1.8% HA1 (equimolar to probucol), or (3) chow plus four-weekly subcutaneous injections of ApoC3 siRNA (5 mg/kg) starting at 6 weeks. ApoC3 siRNA was generously provided by Arrowhead Pharmaceuticals Ltd. All mice had unrestricted access to food and water.

### 2.2. Open Field Test

Anxiety-like behavior was evaluated using the open field. Mice were first acclimated in the dimly lit testing room for 20 minutes, then placed in a square arena (45 × 45 cm, 40 cm high) where they freely explored for 5 minutes. The time each mouse spent in the arena’s periphery was recorded to assess anxiety-like behavior.

### 2.3. Cognitive assessment

In the NOR test, mice were familiarized with two identical objects and assessed for preference toward a novel object 2 hours later to calculate the Preference Index. In the passive avoidance test, mice were trained to avoid the dark compartment in a two-compartment apparatus by receiving a foot shock upon entry and were tested 24 hours later to evaluate avoidance behavior based on latency changes. Detailed methods are provided in the supplementary material.

### 2.4. Sample collection

The mice were euthanised via cardiac puncture under isoflurane anaesthesia. Blood samples were collected into EDTA-coated tubes and plasma was obtained by centrifugation at 3,000 rpm for 10 min at 4 °C and stored at -80 °C. A 2 cm segment of the small intestine duodenum at the proximal end was isolated and fixed in 4% paraformaldehyde The tissues were then processed and embedded in paraffin wax. The brain was carefully removed. The left hemisphere was immediately snap frozen in liquid nitrogen. The right hemisphere was fixed in 4% paraformaldehyde for 24 h and then cryoprotected in 20% sucrose for 72 h at 4°C. The tissues were then frozen in isopentane and stored at -80°C.

### 2.5. Measurement of plasma glucose and insulin

Plasma glucose level was determined using colorimetric assay kit (Caymen Chemical) and plasma insulin concentration was measured using Mouse Insulin ELISA kit (Mercodia) according to the instructions provide by the manufacturers. The optical densities were recorded with the EnSight Multimode Plate Reader, using wavelengths of 500 nm for glucose and 450 nm for insulin.

### 2.6. Measurement of plasma cholesterol, triglyceride and total ApoB

Cholesterol and triglycerides were measured using Randox colorimetric kits (Randox Laboratories, UK) per the manufacturer’s protocol. Briefly, 2 µl of standards or plasma samples were mixed with 200 µl reaction solution in 96-well plates and incubated at 37 °C for 5 minutes. Absorbance was read at 550 nm, and lipid concentrations were determined from the standard curve. Plasma total ApoB was quantified with an ELISA kit (Abcam) following the manufacturer’s instructions. Samples from db/+ mice were diluted 1:3000, while db/db and treated db/db samples were diluted 1:6000 using the provided dilution buffer. Optical density was measured at 450 nm, 15 minutes after adding the stop solution.

### 2.7. Immunofluorescent detection of enterocytic Aβ and ApoB

Intestinal Aβ and ApoB expression were quantified using immunofluorescent microscopy as previously described [4,15]. Paraffin-embedded sections (5 µm) were deparaffinised, rehydrated, and underwent antigen retrieval in boiling deionised water for 15 minutes. Tissues were permeabilised with PBS containing Tween-20 for 10 minutes and blocked with 20% goat serum for 1 hour at 20 °C. Sections were then incubated overnight at 4 °C with rabbit anti-mouse Aβ(1–40/42) antiserum (1:200, Millipore) or polyclonal rabbit anti-mouse ApoB (1:200, Abcam) in blocking solution. Immunofluorescence was visualised using anti-rabbit IgG Alexa546 (1:100, ThermoFisher), and nuclei were stained with HOECHST (1:2000, ThermoFisher). Fluorescent images were captured with a Zeiss Axioscan Z.1 slide scanner and analyzed using Zeiss Zen Blue for pixel intensity in enterocytes.

### 2.8. Plasma Aβ analysis

Plasma levels of Aβ40, Aβ42 and Aβ oligomer were measured using commercially available ELISA kit (Wako) following the manufacturer’s protocol. Their concentrations were determined against the standards of Aβ40 ranging 0–100 pmol/L, Aβ42 ranging 0–20 pmol/L and Aβ oligomer 0-100pmol/L.

### 2.9. Assessment of BBB integrity and tight junction protein and pericyte expression

BBB integrity was evaluated by measuring extravasated plasma IgG in the cortex and hippocampus as previously described [26,27]. Coronal brain sections (20 µm) from snap-frozen left hemispheres were fixed with 4% PFA for 10 min, blocked with 10% donkey serum for 1 h at 20 °C, and incubated overnight at 4 °C with goat anti-laminin a4 (1:200, RnD Systems) in ASEB buffer. Sections were then labelled with donkey anti-goat Alexa555 (1:500, Abcam) for 2 h at 20 °C, followed by anti-mouse or anti-rat Alexa488 for 2 h. Nuclei were stained with HOECHST (1:500, ThermoFisher).

For the immunofluorescence detection of ZO1 and aoccludin-1, 20-μm sections were fixed with 4% PFA followed by incubation with ice-cold acetone/methanol (50:50). Sections were then blocked with 10% donkey serum and incubated overnight with goat anti-laminin a4 and rabbit anti-occludin or anti-ZO-1 (1:500, Thermo Fisher). The sections were incubated with a combination of then incubated with goat a-laminin a4 (1:200, RnD Systems) and either rabbit anti-occludin (1:500, Thermo Fisher Scientific) or rabbit anti-ZO-1 (1:500, Thermo Fisher Scientific) overnight at 4 °C. Pericytes were measured using similar protocol with PDGFRβ (1:100, Cell Signaling Technology). Fluorescent images were captured at 20× on a Zeiss Axioscan Z1 and Zeiss ZEN Intellisis trainable segmentation module was used to identify the pixel intensity of occludin-1 and ZO-1 within vessel area.

### 2.10. Measurement of neuroinflammation, oxidative stress, and axonal and neuronal density

Astrocyte and microglia activation, oxidative stress, and neuronal/axonal density were assessed in the cortex and hippocampus as previously described [14]. Sections (20 µm) from the right hemisphere were rehydrated in PBS, blocked with 10% donkey serum (ThermoFisher) for 1 h at RT, and incubated overnight at 4 °C with the following primary antibodies in ASEB: goat anti-GFAP (1:500, Abcam) for astrocytes, rabbit anti-Iba-1 (1:200, Novachem) for microglia, mouse anti-8-Hydroxyguanosine (8-dOHG) (1:500, Abcam) for DNA oxidation, SMI312 (1:500, Biolegend) for axon density, and NeuN (1:500, Merck) for neuron density. Secondary antibodies included donkey anti-rabbit Alexa488 (1:500, ThermoFisher), donkey anti-goat Alexa555 (1:1000, ThermoFisher), and donkey anti-mouse Alexa647 (1:500, ThermoFisher). Nuclei were stained with HOECHST (1:500, ThermoFisher). Fluorescent images were captured at 20× on a Zeiss Axioscan Z1 slide scanner (Carl Zeiss, Germany) and analysed with Zeiss ZEN Intellisys for cell identification.

### 2.11. Statistical Analysis

All data are expressed as mean ± SEM and were analysed using GraphPad Prism 10. Normally distributed data was analysed using a one-way ANOVA, followed by an uncorrected Tukey’s post hoc multiple comparison test. Non-normally distributed data were analysed by the nonparametric Krusal– Wallis test, followed by Dunn multiple comparisons test. Spearman’s correlation coefficient was used to assess statistical associations. Statistical significance was determined at p < 0.05, with significance levels of P < 0.01, P < 0.001, and P < 0.0001 also reported where applicable.

## 3. Results

### 3.1. Effects of probucol, HA1 and ApoC3 siRNA on metabolic parameters

The db/db mice exhibited significantly elevated plasma glucose and insulin compared with age-matched db/+ controls (Fig. 1A and B), with more profound hyperglycaemia at 28 weeks compared to 14 weeks. The mice were normo-triglyceridaemic but did develop hypercholesterolemia by 28 weeks of age (Fig. 1C and D).

**Figure 1.**
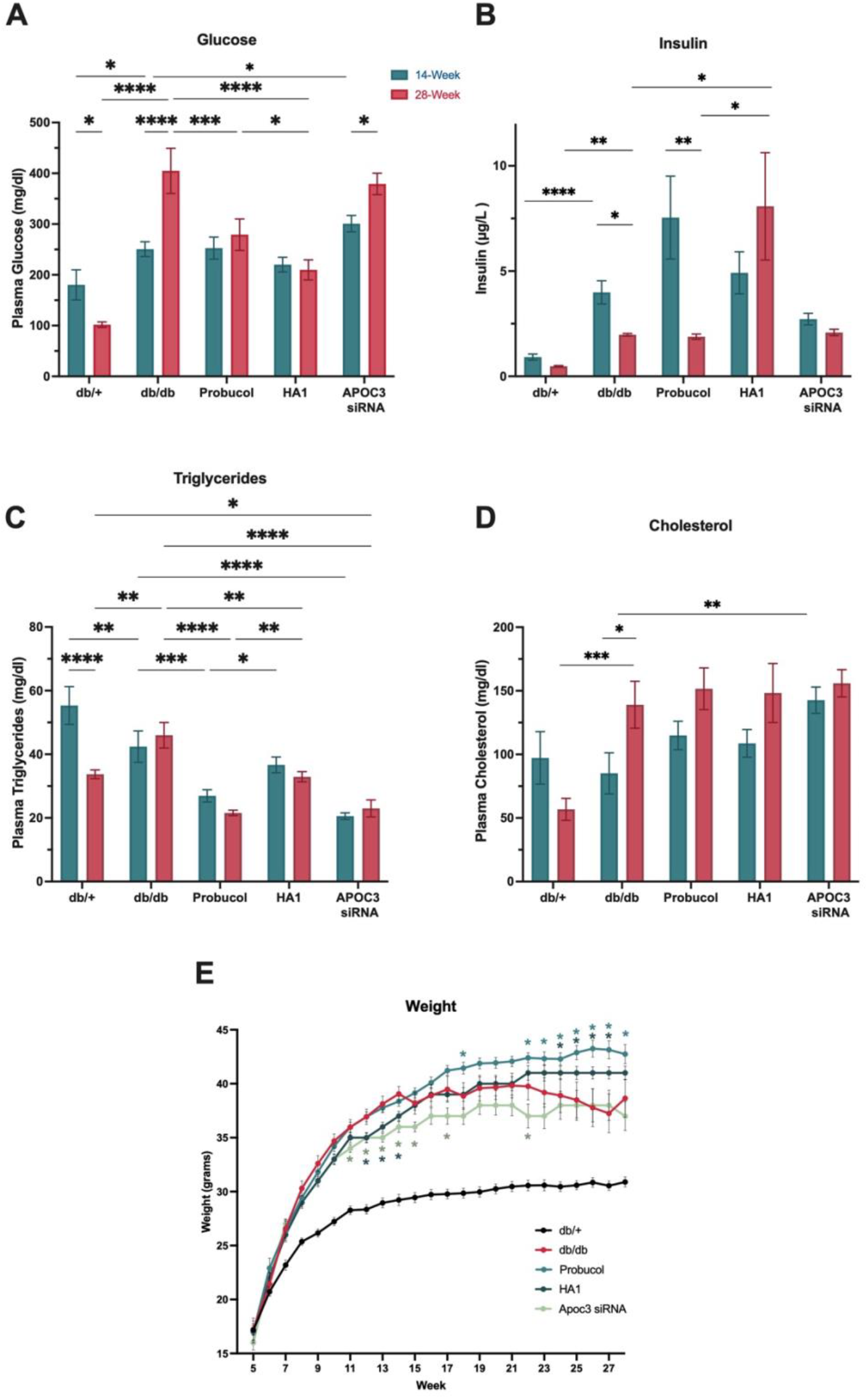
Measurement of metabolic changes in 14- and 28-week-old group. **(A)** non-fasting plasma glucose, **(B)** non-fasting plasma insulin, **(C)** plasma triglycerides, **(D)** plasma cholesterol, **(E)** weight. Values are expressed as ± SEM (7-12 mice per group). Statistical significance was estimated by one-way ANOVA followed by Fisher’s LSD post-hoc test for the data set of glucose, triglyceride, cholesterol, and weight and Kruskal–Wallis for the data set of insulin (n=12-15, *p < 0.05, **p < 0.01, ***p < 0.001, ****p < 0.0001).

Treatment with probucol or HA1 normalised hyperglycemia and exhibited differential effects on plasma insulin homeostasis (Fig. 1A and B). Probucol had no significant effect on plasma insulin compared to control db/+ mice at any age, whereas HA1 increased insulin at 28 weeks. Both probucol and HA1 reduced triglycerides (Fig. 1C). Treatment with probucol and HA1 prevented the weight loss observed in db/db mice. Furthermore, HA1-treated mice exhibited reduced weight gain compared to control db/db mice and probucol-treated mice at 14 and 28 weeks respectively. (Fig E).

Treatment with ApoC3 siRNA significantly increased glucose and cholesterol levels at 14 weeks and led to a marked decrease in triglycerides at both 14 and 28 weeks (Fig. 1A,C and D). No effects on insulin levels were observed in either age group (Fig 1B). Addionally, ApoC3 siRNA-treated mice gained less weight than control db/db mice. (Fig E).

### 3.2. Probucol and HA1 lowered the increase in enterocytic Aβ expression in db/db mice

Enterocytic abundance of Aβ serves as a surrogate marker for amyloid secretion with nascent lipoproteins. The db/db mice had significantly increased enterocytic Aβ at 28 weeks (Fig. 2A and C) which was markedly reduced by probucol and HA1. Conversely, ApoC3 siRNA showed no significant effect (Fig. 2A and 2C).

In db/db mice, exaggerated chylomicron secretion per se is suggested by a significant increase in enterocytic ApoB, an obligatory structural moiety of nascent chylomicrons (Fig 2B and C). Probucol and ApoC3 siRNA had no significant effect on enterocytic ApoB abundance; however, HA1 suggested a transient suppressive effect at 14 weeks, not indicated at 28 weeks (Fig. 2B and C).

**Figure 2.**
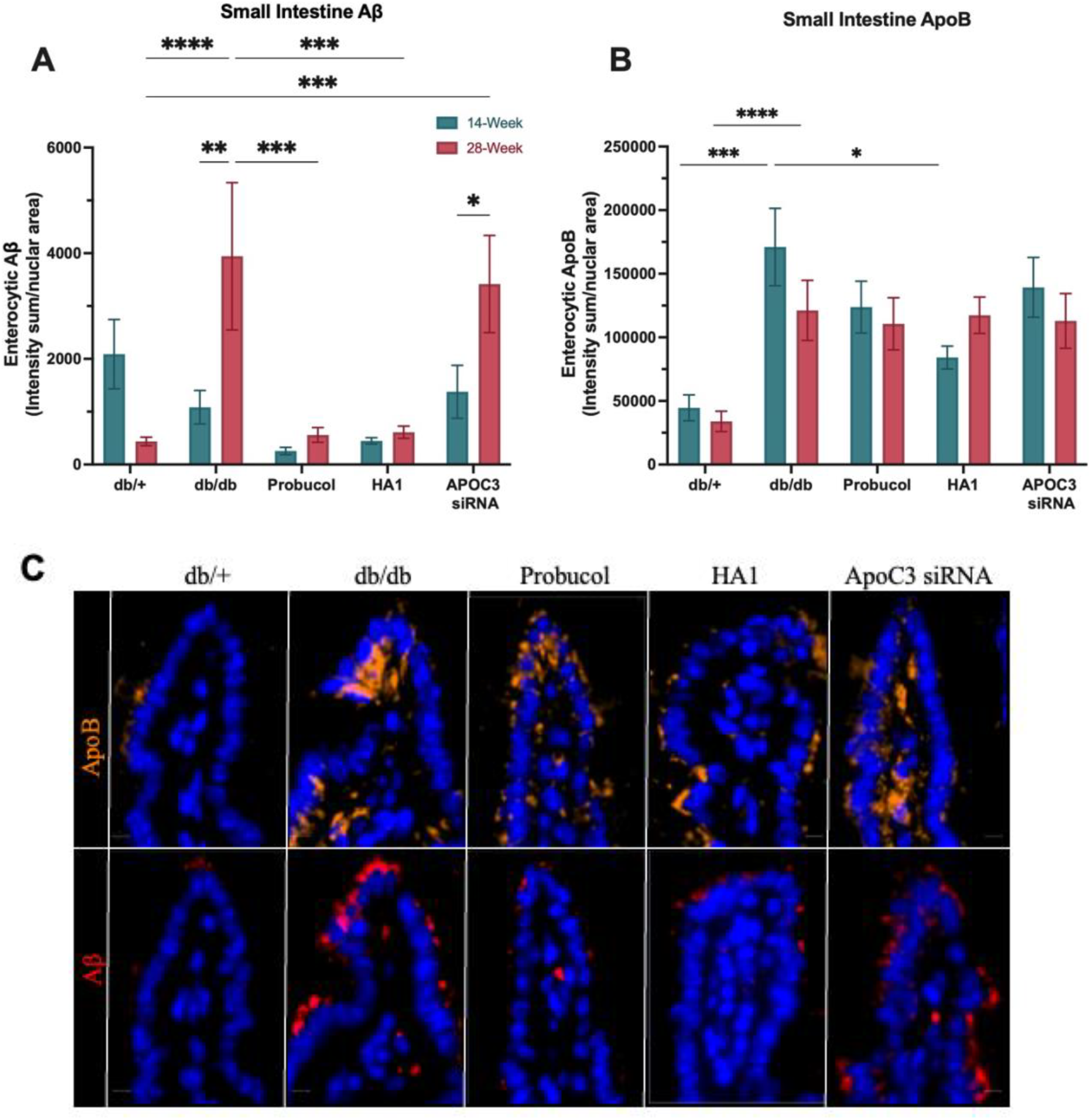
Semi-quantitative immunomicroscopy for enterocytic abundance of Aß and ApoB and correlation coefficient analyses of enterocytic Ab with enterocyte and plasma levels of ApoB. **(A)** the intensity sum of enterocytic Ab expressed per nuclei area, **(B)** the intensity sum of enterocytic ApoB expressed per nuclei area, **(C)** immunofluorescent micrographs of ApoB shown in orange and Aβ shown in red. Statistical significance was estimated by one-way ANOVA followed by Fisher’s LSD post-hoc test for the data set of ApoB, and Kruskal– Wallis for the data set of Aβ (n=12-15, *p < 0.05, **p < 0.01, ***p < 0.001, ****p < 0.0001).

### 3.3. HA1 and ApoC3 siRNA reduced plasma Aβ42 and oligomeric Aβ levels

The more toxic isoform, Aβ_42_, was significantly elevated in the plasma of db/db mice compared to db/+ controls at both ages (Fig. 3A). Additionally, soluble oligomers of Aβ, proposed as potential surrogate markers for amyloid fibril and plaque formation risk, was also increased by approximately 40% in db/db mice at 28 weeks of age (Fig. 3C). Treatment with probucol resulted in a minor, but non-significant, reduction in both Aβ_42_ and Aβ oligomers at 28 weeks. In contrast, HA1 and ApoC3 siRNA treatments significantly lowered these measures at both ages (Fig. 3A). Furthermore, the mechanisms underlying these reductions appear to differ. Probucol and ApoC3 siRNA markedly decreased the plasma abundance of ApoB-containing lipoproteins, which are responsible for chaperoning the majority of plasma amyloid. Conversely, HA1 treatment did not significantly affect plasma ApoB levels (Fig. 3D), consistent with suppression of secretion being the primary mechanism.

**Figure 3.**
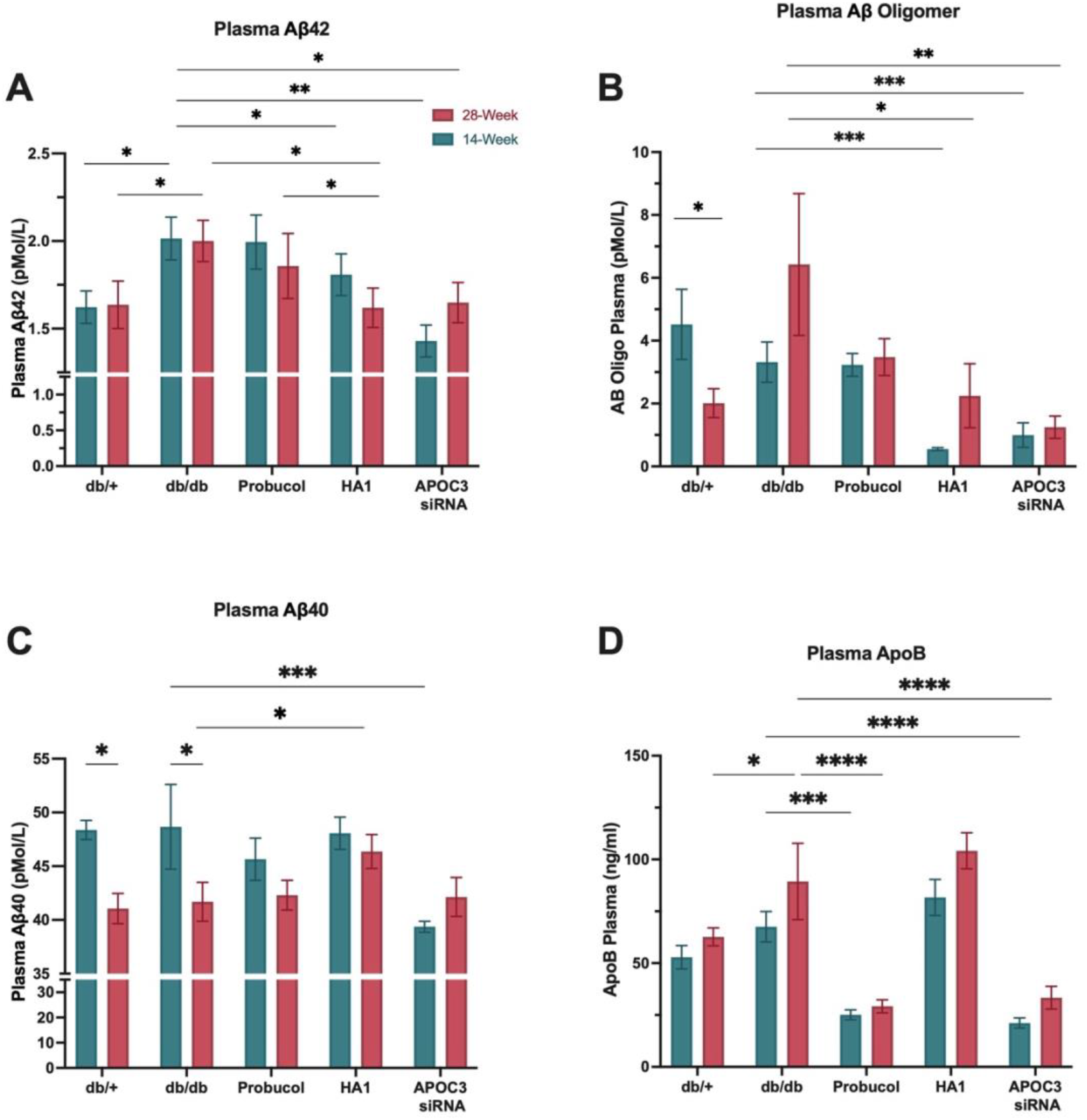
Concentrations of plasma Aβ and ApoB. Pairwise comparisons of plasma (**A**) Aβ42, (**B**) Aβ oligomer, (**C**) Aβ40, and (**D**) plasma ApoB. Values are expressed as ± SEM (9-14 mice per group). Statistical significance was estimated by One-Way ANOVA for the data sets of Ab40, Ab42 and Aβ42/Aβ40 and Kruskal–Wallis for the data set of Ab oligomer ((n=12-15, *p < 0.05, **p < 0.01, ***p < 0.001).

In contrast to Aβ_42_, diabetes did not exert a remarkable effect on the more abundant isoform, Aβ_40_ (Fig. 3B). A weak and transient reduction in plasma Aβ_40_ levels was observed solely with ApoC3 siRNA at 14 weeks (Fig. 3B). No significant effects on Aβ_40_ levels were observed with probucol treatment.

### 3.4. Intestinal expression of chylomicron-Aβ correlates with plasma lipoprotein-Aβ42

Figure 4 examines the relationship between enterocytic Aβ abundance, as a marker of secretion, and plasma Aβ42 levels using data from 28-week-old control, diabetic (db/db), and diabetes-treated groups. Spearman rank correlation analysis revealed a strong correlation between enterocytic Aβ and enterocytic ApoB, suggesting increased synthesis and secretion of chylomicron-Aβ in diabetes. Consistent with the hypothesis that chylomicron kinetics significantly modulate plasma amyloid homeostasis, a positive correlation between plasma Aβ42 and plasma ApoB, as well as between enterocytic Aβ and plasma Aβ42, was observed.

**Figure 4.**
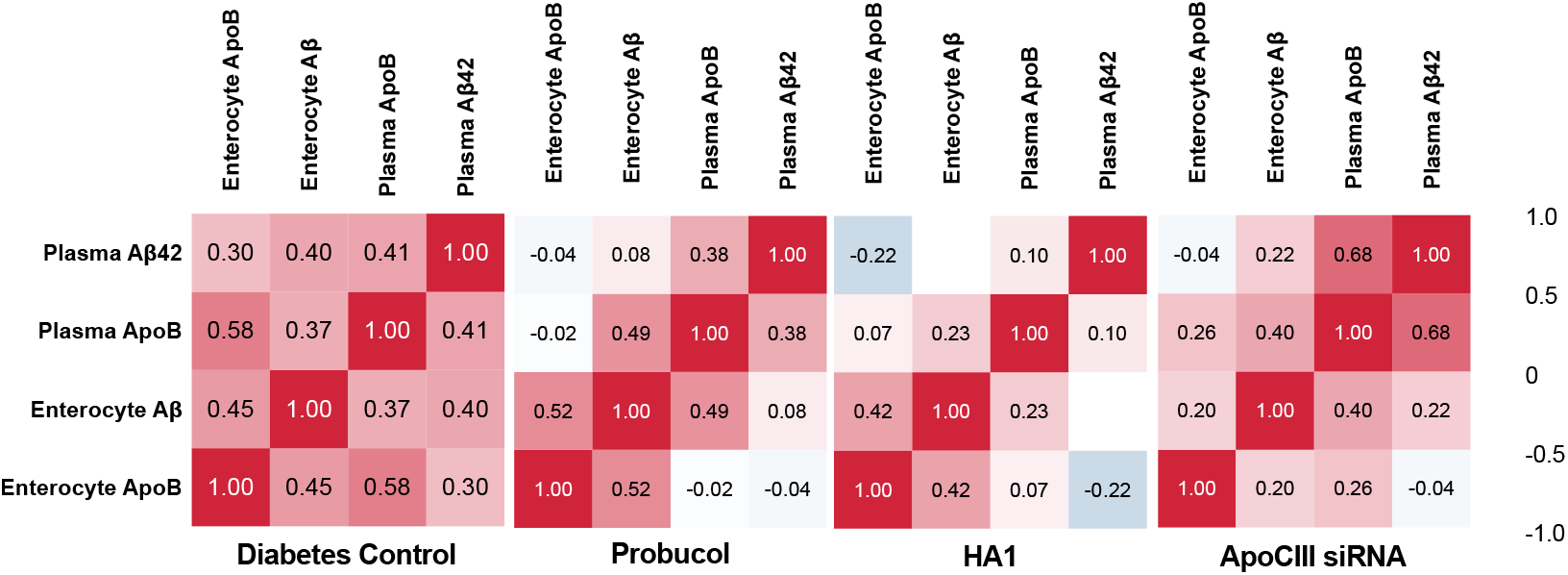
Correlation coefficient analysis between enterocytic Aβ and plasma Aβ42 with lipid parameters. The correlation between enterocytic Aβ and plasma Aβ42 versus enterocytic ApoB, plasma ApoB were considered using Spearman’s correlation coefficient with the data sets of 28-week-old db/+ and db/db mice from the control group and 28-week-old control diabetic and 28-week-old diabetic mice treated with either ApoC3 siRNA, probucol, or HA1 (n=18-24).

In db/db mice treated with probucol, the associations between enterocytic Aβ and enterocytic ApoB, and between plasma Aβ42 and plasma ApoB, were maintained, reiterating the importance of lipoprotein-amyloid assembly and metabolism post-secretion. However, enterocytic Aβ abundance (secretion) no longer correlated with plasma Aβ42 levels. This finding suggests that, in db/db mice, probucol likely influences plasma Aβ42 homeostasis independently of changes in synthesis and secretion.

HA1-treated mice showed a similarly strong correlation between enterocytic Aβ and enterocytic ApoB. However, the association between enterocytic Aβ and plasma Aβ42 was completely abolished. Given that the association between plasma Aβ42 and plasma ApoB was significantly attenuated, these findings suggest that HA1 reduces plasma Aβ42 homeostasis, primarily by selectively inhibiting secretion.

In db/db mice treated with ApoC3 siRNA, the association between enterocytic Aβ and enterocytic ApoB was significantly diminished, suggesting disruption in amyloid synthesis and secretion with nascent lipoproteins. The strong association between plasma Aβ42 and plasma total ApoB indicates that post-secretion metabolism was not significantly impacted.

### 3.5. HA1 increased the expression of tight junction protein ZO1 and pericytes, and effectively mitigated BBB dysfunction

Diabetes markedly exacerbated an age-associated increase in BBB permeability, indicated by the parenchymal abundance of IgG. This effect was indicated in both the hippocampal formation as well as cortex (Fig. 5A). HA1 profoundly suppressed the age- and diabetes-associated increase in parenchymal IgG, concomitant with an increase in tight junction proteins (Fig. 5A-C and E). In db/db mice, probucol did not show efficacy in supporting barrier integrity, with no change observed in tight junction protein expression. Although, ApoC3 siRNA increased the expression of ZO1, it did not prevent IgG leakage. (Fig. 5A-C and E).

**Figure 5.**
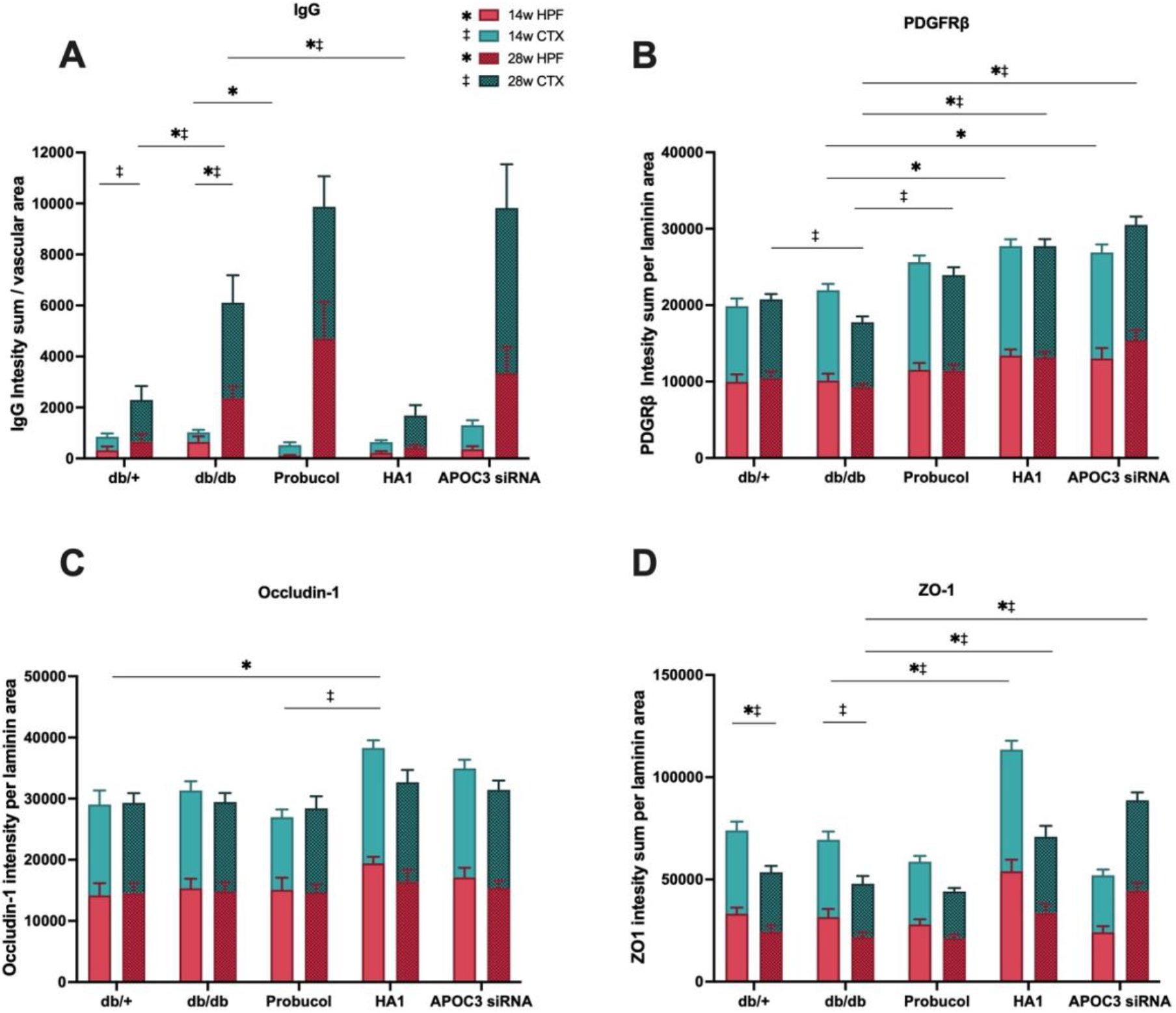
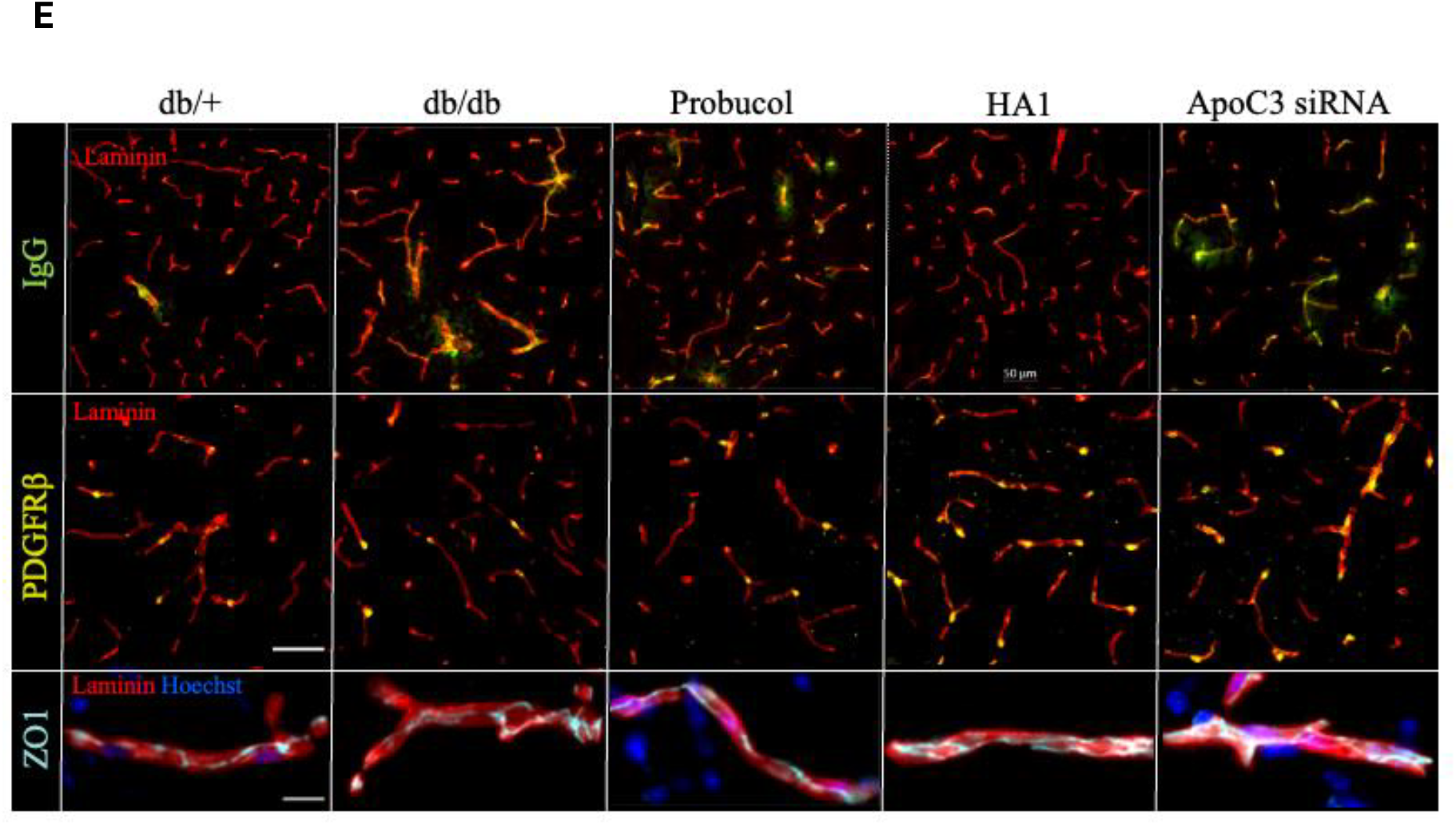
Semi-quantitative immunomicroscopic analysis of markers of cerebral capillary integrity and pericyte and tight junction protein immunoreactivity. (**A**) Hippocampal (HPF) and cortical (CTX) extravasation of plasma IgG expressed as IgG intensity sum per laminin area, (**B**) PDGFRβ expression expressed as intensity sum per laminin area, (**C**) expression of BBB tight junction protein occludin-1, and (**D**) ZO-1 expressed as intensity sum per laminin area. Representative immunomicrographs of IgG (green), PDGFRβ (yellow), and ZO-1 (aquamarine) (**E**). Data are expressed as mean ±SEM. Statistical significance was estimated by one-way ANOVA followed by Fisher’s LSD post-hoc test for the data sets of PDGFRβ, ZO1 and Occludin-1 and Kruskal–Wallis for the data set of IgG ((n=12-15, p<0.05, *significance in hippocampus, †significance in cortex compared to age-matched control).

Pericyte loss/dysfunction is considered as a potential mechanism for diabetes-associated capillary dysfunction. In this study, cortical PDGFRβ levels in db/db mice were significantly lower compared to db/+ mice at 28 weeks. Probucol and particularly, HA1 and ApoC3 siRNA all increased PDGFRβ expression (Fig. 5D and E).

### 3.6. HA1 restored astrocyte levels and demonstrated moderate antioxidative and anti-inflammatory effects in db/db mice

Both 14- and 28-week-old diabetic control mice exhibited a significant reduction in hippocampal GFAP expression compared to the age-matched db/+ controls (Fig. 6A). Astrogliosis was normalised with HA1 treatment and partially restored by probucol at 28 weeks. In contrast, ApoC3 siRNA had no significant effect on GFAP abundance (Fig. 6A and D).

**Figure 6.**
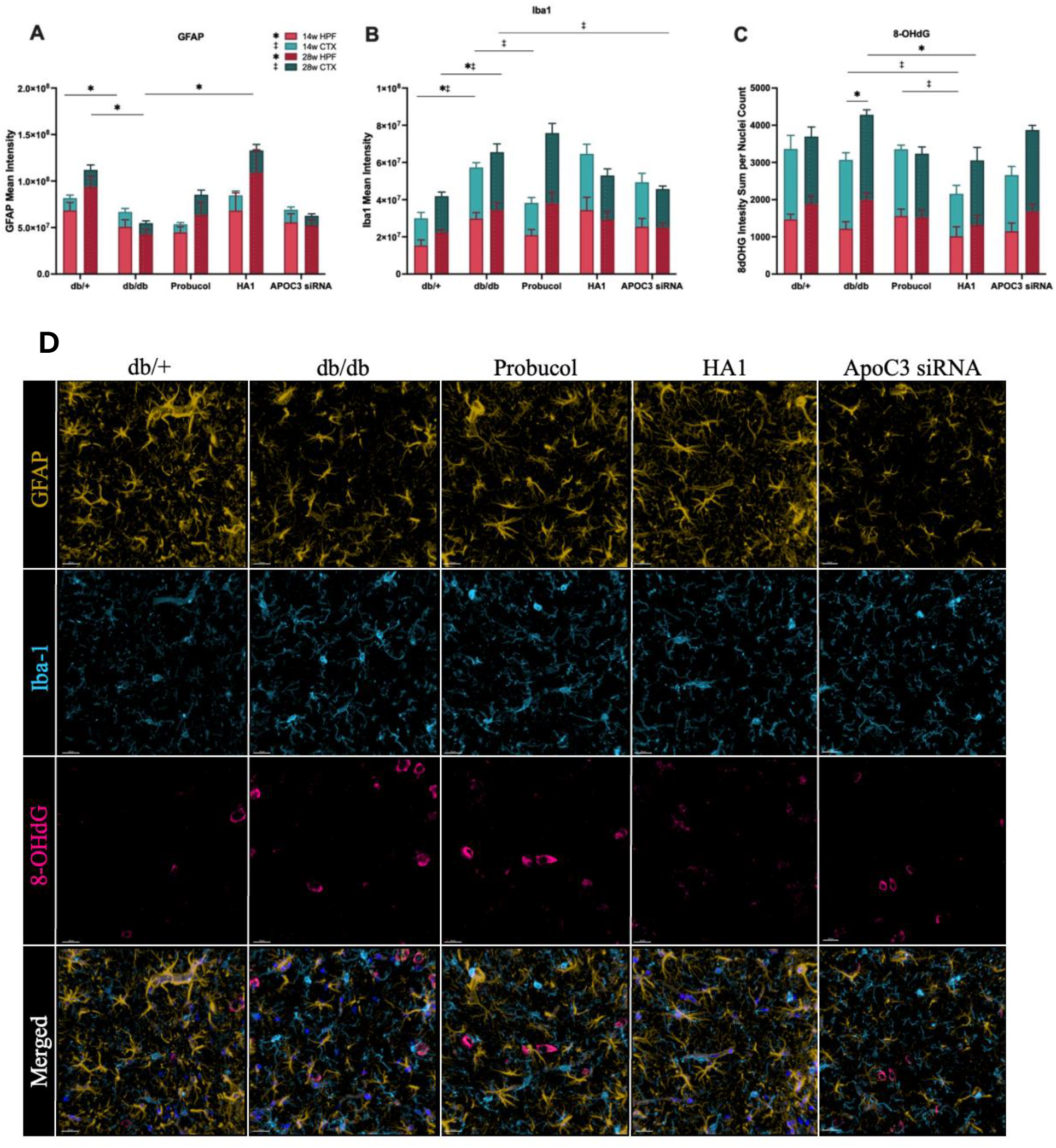
Assessment of oxidative stress and neurovascular inflammation. (**A**) The mean intensity of GFAP in the cortex and hippocampus region, (**B**) the mean intensity of Iba-1 in the cortex and hippocampus region, (**C**) the mean intensity of 8-OHdG in the cortex and hippocampus region, (**D**) representative immunomicrographs of hippocampal GFAP (yellow), Iba1 (blue), 8-dOHG (pink). Gamma settings were adjusted to enhance visualisation: GFAP at 2.0 and Iba1 at 1.5. Data are expressed as mean ± SEM. Statistical significance was estimated by one-way ANOVA followed by Fisher’s LSD post hoc test (n=10-12, p p<0.05, *significance in hippocampus, †significance in cortex compared to age-matched control).

Heightened inflammation is suggested in db/db mice by an exaggerated abundance of Iba1. Probucol transiently delayed the exaggerated Iba1 signature at 14 weeks, whereas ApoC3 siRNA prevented the increase at 28 weeks. HA1 showed minor but non-significant effect at 28 weeks. (Fig. 6B and D).

Downstream of inflammation, the surrogate marker of oxidative stress, 8-OHdG, was also found to be elevated in diabetic conditions but was effectively attenuated by HA1, and to a lesser extent, by probucol and ApoC3 siRNA (Fig. 6C and D).

### 3.7. HA1 and ApoC3 siRNA improved anxiety-like behaviour in 28-week-old diabetic mice

The risk for cognitive decline is markedly increased in T2D and is often preceded by heightened anxiety and behavioural disturbances [16]. At 28 weeks, diabetic mice exhibited heightened anxiety, as assessed by the Open Field Test (Fig 7C) however showed no cognitive deficits in short-term (Novel Object Recognition Test) or long-term memory (Passive Avoidance Test) assessments. (Fig 7A and B). Nonetheless, the intervention with HA1 significantly reduced anxiety-like behaviour in the Open Field Test. Similarly, ApoC3 siRNA normalised behaviour whereas probucol had weaker effects (Fig 7C).

### 3.8. Plasma levels of Aβ correlate with changes in neurovascular unit and anxiety-like behaviour

Spearman’s correlation analysis, shown in Figure 8, explores the association between plasma levels of lipoprotein-Aβ42 and changes in the neurovascular unit, neuroinflammation, oxidative stress, and anxiety-like behavior, based on data at 28 weeks of age (Fig. 8).

**Figure 7.**
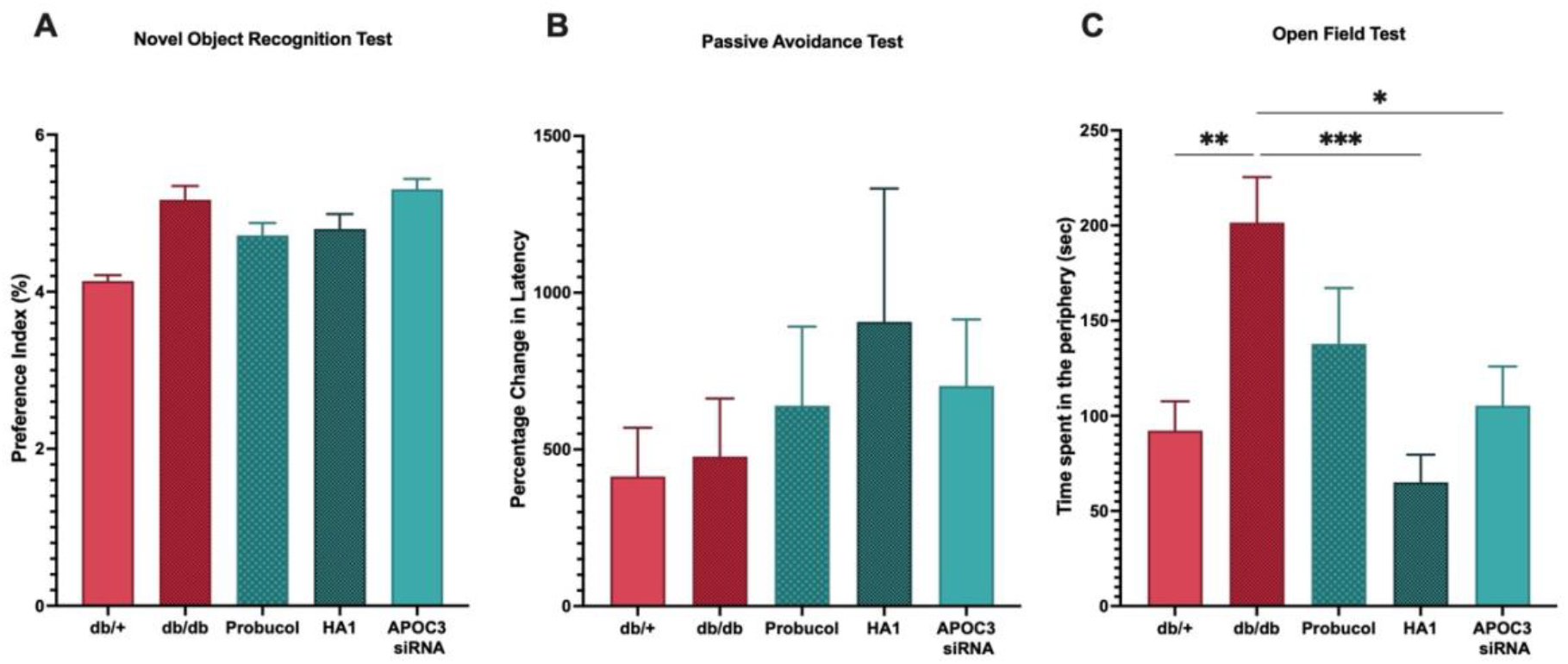
Assessment of anxiety-like behaviour and cognitive function. (**A**) Novel Object Recognition Test, (**B**) Passive Avoidance Test, and (**C**) Open Field Test. Data are expressed as mean ±SEM. Statistical significance was estimated by one-way ANOVA followed by Fisher’s LSD post-hoc test for the data set of NOR and OFT, and Kruskal–Wallis for the data set of PAT ((n=10-15, *p < 0.05, **p < 0.01, ***p < 0.001).

**Figure 8.**
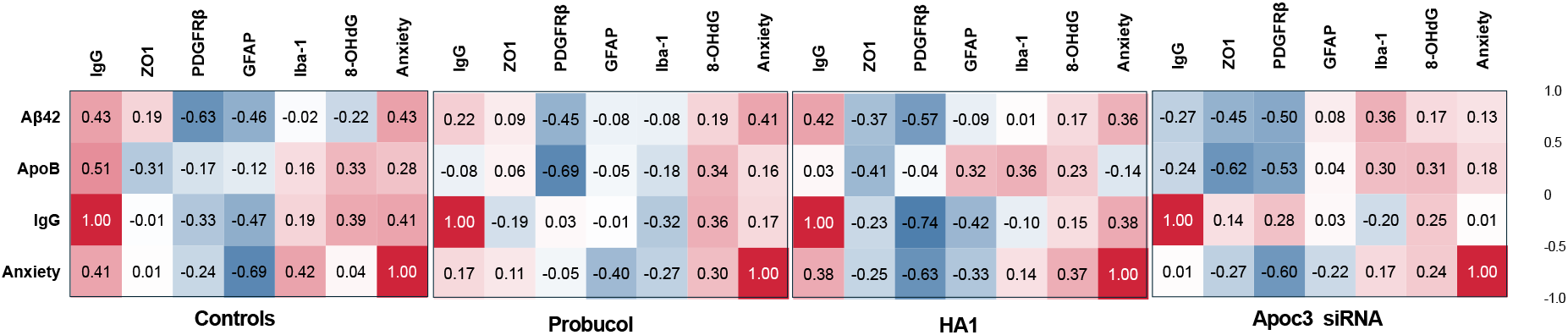
Correlation coefficient analysis between enterocytic Aβ and plasma Aβ42 versus markers of BBB integrity. The correlation between Enterocytic Aβ and plasma Aβ42 versus IgG, GFAP, PDGFRβ and ZO1 were considered with Spearman’s correlation coefficient using the data sets of 28-week db/+ and db/db mice in control group and 28-week-old control diabetic and 28-week-old diabetic mice treated with either ApoC3 siRNA, probucol, or HA1 (n = 18-24).

In control diabetic mice, a positive correlation was observed between plasma lipoprotein-Aβ42 and both IgG leakage and anxiety-like behaviors. Specifically, plasma Aβ42 and BBB leakage were inversely correlated with the homeostasis of astrocytes (GFAP) and pericytes (PDGFRβ), while BBB leakage was positively associated with oxidative stress markers. Furthermore, anxiety-like behavior was significantly associated with elevated plasma lipoprotein-Aβ42, increased BBB leakage, reduced astrocyte (GFAP) levels, and heightened neuroinflammation (Iba-1).

The primary impact of probucol treatment, as suggested by the correlation analysis, was a strong negative correlation between lipoprotein (ApoB), and to a lesser extent plasma Aβ42, with pericyte (PDGFRβ) abundance, though it remains unclear whether these are causally associated. Similarly, probucol treatment showed a relatively strong negative association between astrogliosis (GFAP) and anxiety.

In contrast, HA1-treated mice exhibited strong evidence supporting the regulatory effects of Aβ42 on BBB integrity and anxiety-like behaviours. Specifically, there were robust positive correlations between plasma Aβ42 levels, parenchymal IgG leakage, and anxiety-like behaviours, alongside negative correlations with pericyte (PDGFRβ) levels. Additionally, astrocyte abundance was negatively associated with both IgG leakage and anxiety-like behaviors, with anxiety-like behaviors also showing a positive association with oxidative stress.

Treatment with ApoC3 siRNA had negligible effects on astrogliosis. Nevertheless, correlation analyses suggested a potential link between the expression of the tight junction protein ZO-1 and pericyte homeostasis with lipoprotein-Aβ42 levels and anxiety-like behaviors.

## 4. Discussion

Accumulating research in AD links BBB disruption with increased vascular exposure to elevated levels of circulating lipoprotein-Aβ. Our study investigated these pathological features in a diabetic context using the db/db mouse model, evaluating whether enhancing TRL catabolism through the administration of probucol, the probucol analogue HA1, or ApoC3-targeted siRNA could improve peripheral lipoprotein-amyloid metabolism and alleviate related cerebrovascular abnormalities.

At 14 and 28 weeks, enterocytic ApoB levels were significantly increased compared to controls, indicating exaggerated chylomicron biogenesis and secretion, consistent with previous diabetes studies [17]. However, enterocytic Aβ levels were significantly elevated only at 28 weeks, suggesting that stimulation of lipoprotein biogenesis alone is not the primary driver of plasma amyloidemia. Instead, this likely reflects the direct effects of progressive insulin resistance in db/db mice.

Plasma analyses demonstrated a specific increase in the plasma abundance of the more toxic Aβ_42_ isoform, rather than Aβ_40_, consistent with previous studies in type 2 diabetic individuals and diet-induced obesity models [18]. The increased plasma abundance of chylomicron-bound Aβ_42_, driven by enhanced basal rates of nascent chylomicron synthesis and likely delayed clearance via high-affinity receptor mechanisms [19], resulted in a greater propensity for Aβ oligomer formation. Collectively, chronically exaggerated vascular exposure to postprandial Aβ_42_ and plasma oligomers may contribute to the evolution of BBB dysfunction and neurovascular inflammation in T2DM.

Consistent with a causal role at 28 weeks of age, db/db mice exhibited significantly compromised BBB integrity, as evidenced by the parenchymal extravasation of IgG. However, the observed increase in capillary permeability was not accompanied by measurable reductions in the expression levels of the tight junction proteins ZO-1 and occludin. In early diabetes, tight junction protein expression may initially increase as a compensatory response to maintain barrier integrity [21], but ultimately fail to sustain proper function due to abnormal localisation [22]. Moreover, BBB integrity is dependent on the structural and functional support provided by pericytes and astrocytes and is particularly vulnerable to oxidative stress and inflammation. In this study, db/db mice exhibited reduced pericyte coverage of cerebral vessels and decreased astrocyte populations, along with elevated markers of oxidative stress and reactive microgliosis.

By 28 weeks of age, the evolution of BBB dysfunction in db/db mice had not progressed to frank neurodegeneration or loss of axonal density (Fig S1.). Cognitive performance also remained unaffected, with no deficits observed in both long-term and short-term memory, as assessed by the PA and NOR tests. These findings align with our previous studies, which indicate that BBB impairment precedes the onset of overt neurodegenerative and cognitive symptoms [23]. However, behavioural analysis using the open field test revealed a significant increase in anxiety-like behaviours in db/db mice. The development of anxiety- and depressive-like phenotypes is considered a prodromal symptom of Alzheimer’s disease (AD) [24], potentially driven by increased permeation of proinflammatory cytokines into the brain parenchyma and activation of cerebral glial cells [25]. Consistent with this, Spearman correlation analysis indicated an association between anxiety-like behaviours and heightened capillary permeability, microglial activation, and astrocyte loss. Additionally, elevated plasma levels of lipoprotein-Aβ_42_ were associated with BBB dysfunction, degeneration of astrocytes and pericytes, as well as increased anxiety-like behaviour. These findings suggest that lipoprotein-Aβ_42_ may compromise BBB permeability and contribute to behavioural deficits by inducing neuroinflammation and altering the expression and function of astrocytes and pericytes. Indeed, previous studies have demonstrated that hippocampal astrocytes can exert anxiolytic effects by enhancing synaptic activity [26], while both astrocytes and pericytes are critical for maintaining BBB integrity and responding to pro-inflammatory stimuli, thus modulating the neuroinflammatory environment [20].

Of the three interventional agents studied—native probucol, HA1, and ApoC3 siRNA, HA1 had the most significant positive effects. It ameliorated hyperglycaemia, lowered plasma triglycerides and improved insulin secretion. HA1 markedly decreased the secretion of chylomicron-Aβ and reduced plasma levels of the toxic lipoprotein-associated Aβ_42_ and Aβ oligomers without altering plasma ApoB concentrations, which aligns with studies suggesting that exposure to lipoprotein-amyloid, rather than ApoB itself, regulates capillary function [21,22]. HA1 also increased tight junction protein expression, maintained astrocyte and pericyte homeostasis and prevented cerebral oxidative stress and microglial activation, leading to a significant reduction in anxiety-like behaviours in db/db mice.

Correlational analyses further revealed that reductions in plasma Aβ_42_ in HA1-treated mice were associated with improved BBB integrity, increased ZO-1 and pericyte levels and reduced anxiety-like behaviour. Additionally, elevated astrocyte and pericyte levels correlated with enhanced BBB integrity and reduced anxiety. These findings indicate that HA1 contributes to the preservation of BBB function, mitigating behavioural deficits by preventing vascular exposure to Aβ_42_ and maintaining neurovascular and glial components.

Previous studies in high-fat-fed models demonstrated that native probucol significantly attenuates diet-induced chylomicron-Aβ secretion, accompanied by notable preservation of the BBB and cognitive function [10,15]. This study extends those findings by exploring probucol’s effects in a robust model of insulin resistance. Probucol improved several metabolic parameters, including plasma glucose, triglycerides, and ApoB concentrations. While probucol demonstrated transient efficacy in maintaining BBB integrity up to 14 weeks of intervention, this effect was not sustained with chronic diabetes at 28 weeks. This contrasts with findings from high-fat-fed mice, where capillary preservation was observed up to 12 months of age[23]. Nonetheless, the present findings align with the concept of a lipoprotein-amyloid/capillary axis in neurodegeneration, as native probucol did not chronically reduce plasma levels of chylomicron-Aβ_42_ or oligomers, nor prevent BBB breakdown. Supporting evidence from Meakin et al. [37] showed that elevated plasma Aβ_42_ levels in high-fat-fed, obese, and hyperglycaemic mice induced vascular dysfunction similar to that observed in mice on a regular chow diet intravenously administered physiological doses of Aβ_42_.

ApoC3 siRNA markedly accelerates the metabolism and clearance of nascent TRLs by promoting endothelial LPL-mediated hydrolysis and conversion to triglyceride-depleted particles, which are primarily cleared via hepatic TRL and remnant receptors [13]. While ApoC3 siRNA has not been shown to inhibit lipoprotein biogenesis, it may increase chylomicron secretion and, by extension, potential lipoprotein-Aβ flux [24]. In line with the proposed mechanisms of accelerated lipoprotein metabolism and clearance, this study demonstrates, for the first time in a murine model of insulin resistance, a significant reduction in plasma levels of ApoB, Aβ_40_, and Aβ_42_, likely reflecting accelerated clearance of lipoprotein-amyloid without significantly altering the secretion of lipoprotein-Aβ (enterocytic abundance). However, ApoC3 siRNA treatment did not confer microvascular protection, despite lower plasma amyloid levels as indicated by static measures. This highlights the need to consider absolute quantitative measures of metabolism (concentration × turnover). Postprandial (chylomicron) lipid flux is orders of magnitude greater than endogenous (VLDL) hepatic lipid flux, which may also apply to lipoprotein-associated amyloid [25].

Another potential confounder to ApoC3 siRNA’s efficacy could be heightened microvascular exposure to non-esterified fatty acids (NEFAs), which are liberated in greater amounts with ApoC3 siRNA treatment due to increased triglyceride lipolysis. Multiple studies have shown that lipolysis products from TRLs can damage endothelial barrier function, induce apoptosis in endothelial cells and elevate BBB permeability in murine models [26,27]. Additionally, NEFAs may exacerbate insulin resistance [28]. Consistent with this, ApoC3 siRNA treatment in db/db mice worsened hyperglycemia and elevated cholesterol levels at 14 weeks. However, in this study db/db mice were not profoundly hypertriglyceridemic.

While neither probucol nor ApoC3 siRNA treatments mitigated capillary leakage, both may have potentially improved some aspects of capillary integrity, which could be linked to reduced anxiety measures. Specifically, attenuation of anxiety was associated with increased pericyte and tight junction protein ZO-1 abundance, along with reduced oxidative stress.

Of the agents studied, HA1 demonstrated superior efficacy compared to parent compound probucol and ApoC3 siRNA in preserving neurovascular integrity and mitigating behavioural deficits in the robust db/db murine model of insulin resistance. The physiological mechanisms suggest that HA1 reduces absolute exposure to the particularly cytotoxic lipoprotein-Aβ42, thereby strengthening barrier properties and synergistically attenuating inflammation. However, this study was not powered to elucidate how HA1 modulates the synthesis and secretion of Aβ, which is secreted as an apoprotein of nascent TRLs. It remains unclear whether, in enterocytes and hepatocytes, Aβ is derived from precursor proteins found in intracellular and plasma membranes. Colocalisation studies of enterocytic Aβ with ApoB suggest that Aβ may originate from enterocytes [29].

A significant consideration is that HA1 is a novel chemical entity conjugated to probucol at the C24 position of the lithocholic acid (LCA) moiety at the phenolic end of probucol. LCA itself may mediate biological effects that synergize with probucol’s pleiotropic properties. LCA can activate the FXR and TGR5 signalling pathways, which induce the release of glucagon-like peptide-1 (GLP-1) [30]. GLP-1 has been shown to reduce food intake, promote weight loss, increase insulin secretion, and suppress lipid synthesis [31]. Moreover, bile acid-mediated FXR activation has been shown to lower postprandial lipemia [32,33] and downregulate MTP expression [34,35]. In this study, HA1 treatment indeed reduced weight gain, lowered glucose and triglyceride levels, and elevated insulin levels. LCA is also a potent activator of vitamin D receptor which may further enhance its protective effects [36].

More broadly, the focus on postprandial (chylomicron) amyloid metabolism, in this study, provides valuable insights into the interplay between dietary lipid processing and endogenous (VLDL) amyloid metabolism in diabetes. After the ingestion of fats, postprandial chylomicrons will significantly influence the circulating lipoprotein-amyloid environment and can modulate the transport and deposition of amyloid peptides [21]. In individuals with diabetes, postprandial aberrations in lipoprotein metabolism often results also in increased production and impaired clearance of VLDL, potentially exacerbating amyloid accumulation in the brain[37]. By understanding how postprandial chylomicrons interact with amyloid metabolism, we aim to elucidate the complex mechanisms that link dietary fat intake, endogenous lipoprotein-amyloid metabolism and amyloid-related pathologies in diabetic conditions. This understanding may inform therapeutic strategies aimed at mitigating cognitive decline associated with these metabolic disturbances.

This study in a preclinical db/db murine model of diabetes has limitations that impact its translational relevance. While the model is useful for studying diabetic metabolic changes, it does not fully replicate human diabetes complexity. Future studies could incorporate additional animal models to enhance applicability to human physiology. Another limitation is the lack of detailed lipoprotein compositional data, which would offer deeper insights into lipid metabolism alterations associated with diabetes. Including high-resolution lipoprotein profiling in future research could clarify biochemical changes linked to disease progression and treatment responses.

In conclusion, this study highlights the complex interplay between lipoprotein-amyloid metabolism, BBB integrity and neuroinflammation in a diabetic context, using the db/db mouse model. The findings reinforce the hypothesis that chronic exposure to lipoprotein-Aβ_42_, exacerbated by dysregulated TRL biogenesis and delayed clearance, significantly compromises neurovascular integrity. Of the therapeutic agents tested, the second-generation probucol analogue HA1 demonstrated superior efficacy in alleviating hyperglycemia, improving metabolic profiles and reducing plasma levels of toxic Aβ_42_ and Aβ oligomers. HA1 treatment preserved BBB integrity by maintaining astrocyte and pericyte homeostasis and reducing oxidative stress and microglial activation, thereby preventing anxiety-like behaviours—an early prodromal marker of AD. In contrast, while native probucol and ApoC3 siRNA showed transient, or partial effects on metabolic parameters and amyloid clearance, they were less effective in long-term neurovascular protection. The multifactorial benefits of HA1, including its potential synergy with bile acid signalling pathways, suggest broader therapeutic applicability beyond diabetes-induced neurovascular pathologies, offering new avenues for the treatment of neurodegenerative diseases characterized by amyloidosis and vascular dysfunction. Further investigation into the precise mechanisms by which HA1 modulates amyloid synthesis and secretion, particularly in the context of enterocytic and hepatic metabolism, will be critical in advancing its therapeutic development.

## Supporting information

Supplemental Analysis

## Author’s contributions

**Arazu Sharif:** Methodology, Formal analysis, Data curation, Conceptualization, Writing – original draft. **John Mamo:** Funding acquisition, Conceptualization. Writing – review & editing, Supervision. **Virginie Lam:** Funding acquisition, Writing – review & editing, Supervision. **Gerald F Watts:** Resources, Conceptualization, review & editing. **Giuseppe Luna:** Methodology. **Michael Nesbit:** Methodology, Formal analysis. **Hani Al-Salami:** Resources, Conceptualization. **Armin Mooranian:** Resources, Conceptualization. **Ryu Takechi:** Methodology, Funding acquisition, Data curation, Conceptualization, Writing – review & editing, Supervision.

## Data and Material Availability

The datasets used in the current study are available from the corresponding author on reasonable request.

