## Supplemental Analysis for "Effects of HA1 (a Probucol Analogue) and ApoC3 siRNA on Lipoprotein-Amyloid Metabolism, Neurovascular Integrity and Cognitive Function in db/db Mice"

**Methods:**

**Novel Object Recognition test**

The Novel Object Recognition (NOR) test was conducted in a dimly lit room, using the HVS Image 2014 software (HVS Image, UK) within a square apparatus measuring 45 cm on each side, with a wall height of 40 cm. The experiment was structured into three phases: habituation, familiarization, and testing. During the habituation phase on the first day, mice were allowed to explore the empty arena for 5 minutes to acclimatise to the environment. 24 h later, during the familiarisation phase, each mouse was reintroduced to the arena where two identical rectangular objects were placed 20 cm apart. The mice were allowed to explore these objects for 10 minutes before being returned to their home cages. Following a 2h delay, the testing phase commenced. Mice were placed back in the arena, where one of the previously introduced rectangular objects was replaced with a curved object of comparable dimensions. The exploration period was set at 5 minutes. The Preference Index (PI) was calculated as the ratio of the time spent exploring the novel object to the total exploration time of both objects, using the formula:

PI = ((Time exploring novel object) / (Time exploring novel object + Time exploring familiar object)) × 100.

For inclusion in the results, mice were required to meet a minimum familiarization threshold of 10 seconds of active exploration per object during the familiarization phase.

**Passive Avoidance test:**

The passive avoidance test was conducted using a two-compartment apparatus consisting a brightly lit compartment (1000 lux) and an  dark compartment (Ugo Basil, Italy). During the training session, mice were placed in the bright compartment for 30 seconds before the door to the dark compartment was opened. Mice had 300 seconds to enter voluntarily. Upon entry, a 0.3 mA foot shock was administered for 2 seconds. Subsequently, mice were left in the dark compartment for 10 seconds before being transferred back to their home cages.

24 h later, the retention test was conducted. Mice were placed in the bright compartment and after a 30-second delay, the door to the dark compartment was opened again. A maximum test period of 315 seconds was allowed for the mice to enter; no shock was administered upon entry during this phase. Avoidance percentage was calculated based on the change in latency times between the training and test phases, using the formula:

Avoidance % = ((Test phase latency - Training phase latency) / Training phase latency) × 100.

**Results:**

To assess whether enhancements in neurovascular integrity positively affected axonal and neuronal density, we quantified SMI312 immunoreactivity and the percentage of NeuN-positive cells within the dentate gyrus of the hippocampus using immunofluorescent staining. The diabetic db/db mice exhibited no changes in axonal and neuronal density compared to db/+ control mice at either 14 or 28 weeks. Treatment with probucol and HA1 increased neuronal density at 14 weeks, but no changes in axonal and neuronal densities were noted in response to apoc3 treatment in either age group.

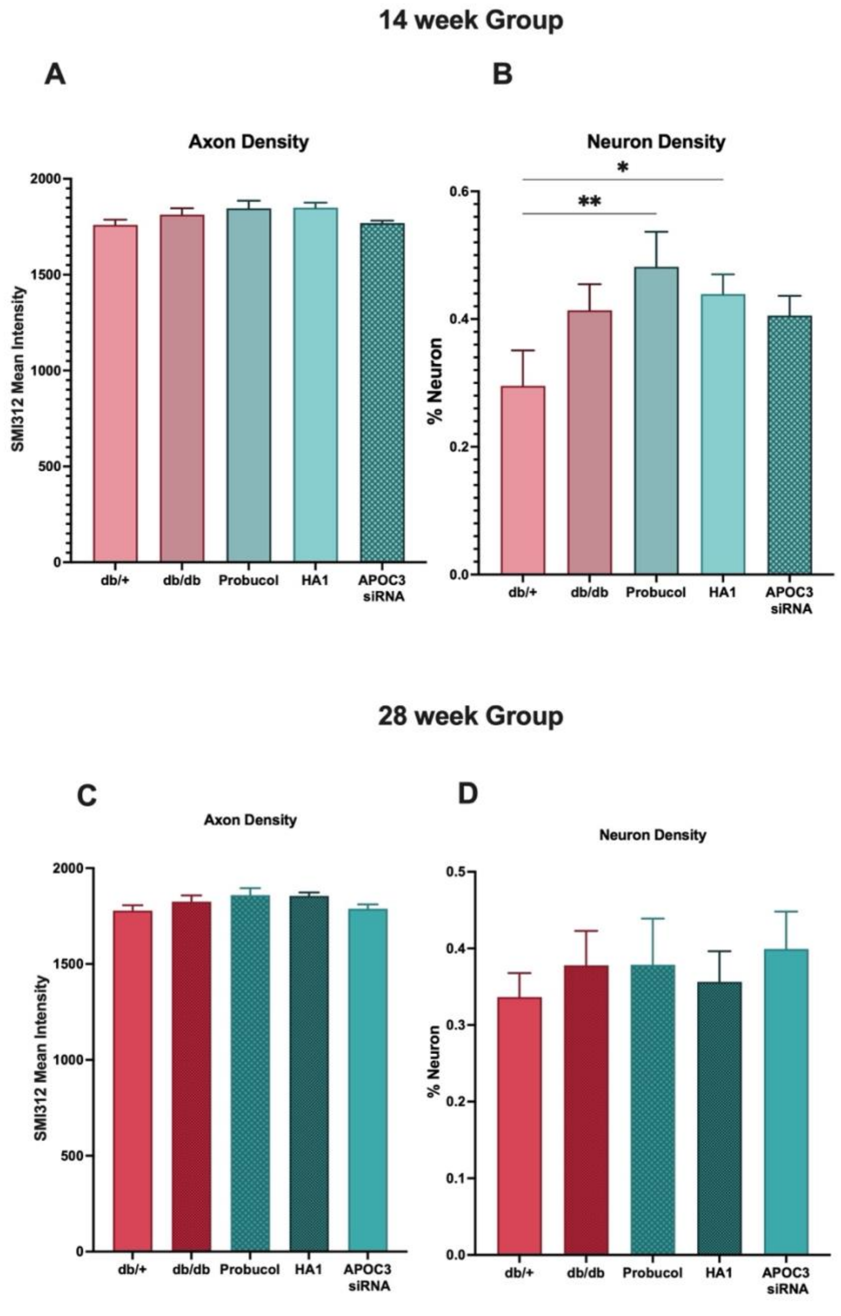


**Figure S1. Assessment of neuron and axon density with the dentate gyrus of hippocampus.** (**A**) Axon and (**B**) neuron density at 14 weeks. (**C**) Axon and (**B**) neuron density at 28 weeks. ±SEM. Statistical significance was estimated by one-way ANOVA followed by Fisher's LSD post-hoc test for the data set of NOR and OFT, and Kruskal–Wallis for the data set of PAT (*p < 0.05, **p < 0.01, ***p < 0.001).
